# Severe Osteoarthritis in Aged PANX3 Knockout Mice: Implications for a Novel Primary Osteoarthritis Model

**DOI:** 10.1101/2023.07.04.547676

**Authors:** Brent Wakefield, Justin Tang, Julián Balanta Melo, Jeffrey L. Hutchinson, Rehanna Kanji, Geneva Herold, Brooke L. O’Donnell, Courtney Brooks, Patti Kiser, Matthew W. Grol, Cheryle A. Séguin, Lilian I. Plotkin, Frank Beier, Silvia Penuela

**Author notes:** Co-first authors (Brent Wakefield, Justin Tang). To whom correspondence should be addressed: Silvia Penuela, Ph.D., Associate Professor, Department of Anatomy and Cell Biology, University of Western Ontario, London, Ontario, Canada.

## Abstract

Osteoarthritis (OA) is a multi-factorial disease associated with aging. As the molecular mechanisms underpinning the pathogenesis of this disease are unclear, there are no disease-modifying drugs to combat OA. Pannexin 3 (PANX3) has been shown to promote cartilage loss during posttraumatic OA. In contrast, the ablation of *Panx3* in male mice results in spontaneous full-thickness cartilage lesions at 24 months of age. While protected from traumatic intervertebral disc (IVD) degeneration, *Panx3* knockout (KO) mice show signs of IVD disease with altered disc mechanics. Whether the deleterious effects of ablating *Panx3* in aging is the result from accumulated mechanical damage is unknown. We used male and female wildtype (WT) and global *Panx3* KO C57Bl6 mice aged to 18 months of age. Mice were then randomized to sedentary (SED) or forced treadmill running (FEX) for 6 weeks. Knee joint tissues including patellar tendon, quadriceps and distal patellar enthesis, and synovium were analyzed histologically and through micro-CT, along with lumbar spine IVDs. Half of male and female sedentary *Panx3* KO mice developed full-thickness cartilage lesions, severe synovitis, and ectopic fibrocartilage deposition and calcification of the knee joints in comparison to all other conditions. *Panx3* KO mice with severe OA show signs of quadriceps and patellar enthesitis, characterized by bone and marrow formation. Forced treadmill running did not seem to exacerbate these phenotypes in male or female *Panx3* KO mice; however, it may have contributed to the development of lateral compartment OA. The IVDs of aged *Panx3* KO mice displayed no apparent differences to control mice, and forced treadmill running had no further effects in either genotype. We conclude that aged *Panx3* KO mice show features of late-stage primary OA including full-thickness cartilage erosion, severe synovitis, and enthesitis. These data suggest that the deletion of *Panx3* is deleterious to synovial joint health in aging.

**LAY SUMMARY:** Osteoarthritis is a common joint disease, and its incidence can increase with aging. As such, no drugs are available to treat its progression. Recently, we discovered that cellular channels composed of Pannexin 3 (PANX3) proteins may contribute to the health of joint tissues. In this study, we propose a new model for age-related primary OA, using histology, microCT, and pathology assessments under sedentary and forced exercise conditions. We found that aged mice lacking the channel protein PANX3 developed late-stage OA, including severe cartilage loss, joint inflammation, and tendon attachment damage. Half of the sedentary PANX3-deficient mice showed these changes, along with abnormal bone growth. Treadmill running did not make the condition worse but may have caused additional joint damage in some areas. The spine, however, remained unaffected. These findings suggest that deleting *Panx3* harms joint health with aging. Importantly, this mouse model mimics human primary OA, even without added factors like diet changes or forced exercise.

## 1.1 Introduction

Globally, knee osteoarthritis (OA) affects millions—23% of the population over 40—causing disability that results in enormous personal and socioeconomic burden [1]. While several risk factors exist for developing OA including sex, obesity, previous joint injury, joint shape and alignment, aging is the single greatest risk factor [2]. The molecular mechanisms that underpin this age-associated destruction of synovial joints are poorly understood.

Pannexin 3 (PANX3) is a channel-forming glycoprotein expressed in osteoblasts where it can act as a Ca^2+^ release channel at the endoplasmic reticulum [3] and in chondrocytes where it can act as an ATP release channel at the cell membrane [4]. PANX3 is also expressed in annulus fibrosus (AF) cells of the intervertebral disc (IVD) [5, 6]. In an early rat model of traumatic OA (anterior crucial ligament transection and partial medial meniscectomy), *Panx3* mRNA expression was upregulated in OA cartilage compared to control knees [7]. Additionally, in a 30-week-old mouse model, PANX3 was upregulated in cartilage following destabilization of the medial meniscus (DMM) surgery to induce post-traumatic OA [8]. This increased expression of PANX3 in diseased cartilage suggests its mechanistic involvement in traumatic OA development in rodent models. In fact, Moon *et al.* performed the DMM surgery on global and chondrocyte specific *Panx3* knockout (KO) mice and found that these mice were strongly protected from OA compared to wildtype (WT) mice [8]. Similarly, in an IVD injury model, *Panx3* KO mice had fewer hypertrophic cells of the AF, and the AF structure was largely preserved compared to WT mice [6]. These two models suggest that the absence of PANX3 is protective in traumatic/injury-induced joint disease. In humans, PANX3 is upregulated in OA cartilage tissue [8], and noncoding intronic single nucleotide polymorphisms (SNPs) of *PANX3* are strongly associated with chronic low back pain [9]. These data suggest that PANX3 function in cartilage is conserved across rodents and humans and may be an important molecular player of OA.

Previous studies have shown that aging influences the genetic expression patterns of joint tissues in response to stress/injury [10–12], and therefore, aged models are required to better understand the pathobiology of age-associated OA. To this point, male *Panx3* KO mice at 18 or 24 months of age showed accelerated cartilage erosion, subchondral sclerosis, and synovitis of the knee joint [13], which was in contrast to the previously seen protective effects in the DMM model in adult mice [8]. An important difference between the two studies is that the aged mice were given a running wheel in their cage for environmental enrichment, which could have contributed to the different effects of *Panx3* KO on joint tissues. In the IVD, we have shown that uninjured IVDs are sensitive to aberrant biomechanical loading [6], again highlighting the context-dependent function of PANX3 in joint health. To gain more insight into how mechanical loading affects joint wear and tear in conjunction with a genotypic deletion of *Panx3*, we recently examined a forced-exercise model on 24-30-week-old with a treadmill running protocol. Consistent with our prior findings, *Panx3* exhibited protective effects on the knee joints, as observed with *Panx3* KO mice exhibiting more pronounced superficial defects in tibial cartilage. Additionally, PANX3 mice displayed greater histological features of IVD degeneration (IDD) after forced treadmill running [14].

In this study, we investigated how aging and excessive mechanical use, via forced treadmill running as applied in the 24–30-week-old mice, influence joint pathology in *Panx3* KO mice. *Panx3* KO mice demonstrated a bimodal distribution in which roughly half of the animals, regardless of forced running, had full-thickness cartilage erosion down to the subchondral bone and expanded synovium containing ectopic calcification and fibrocartilage with scattered lymphocytes, while the other half had mild synovitis. In contrast, all WT mice had mild signs of superficial cartilage erosion and synovitis under either treatment. Male *Panx3* KO mice also displayed cartilage, bone, and bone marrow in the quadriceps enthesis reminiscent of enthesitis. The degree to which these mice had enthesitis or tendinopathy of the knee was correlated with OA scores. At the lumbar spine, *Panx3* KO mice IVDs were histologically similar to WT mice. These results suggest that aged *Panx3* KO mice, regardless of sex and activity, develop severe age-related knee joint pathology including OA in aging.

## 1.1 Methods

### 1.1.1 Mice

Animals used in this study were bred in-house and euthanized in accordance with the ethics guidelines of the Canadian Council for Animal Care. Animal use protocols were approved by the Council for Animal Care at Western University Canada (AUP 2019-069). C57BL/6N Mice were housed in standard shoe box-style caging, exposed to a 12-hour light/dark cycle, and ate regular chow *ad libitum*. WT and *Panx3* KO mice were congenic [8]. DNA was collected from ear clippings of each mouse to determine the genotype using polymerase chain reaction (PCR) as previously described [8, 15]. At sacrifice, mouse knees and spines were collected and immediately processed for histological analysis.

### 1.1.2 Forced treadmill running

At 18 months, mice were randomized to either a no exercise group (sedentary, SED) or a forced treadmill running (forced-exercise, FEX) group. FEX groups ran on a treadmill (Columbus Instruments, Ohio) for 6 weeks, 1 hour a day for 5 days a week, at a speed of 11LJm/min, and a 10° incline—an adapted protocol that has previously been used to induce OA in male C57BL/6 mice [16]. All mice were continuously monitored by an operator in the room to ensure completion of the running protocol. Additionally, the mice were encouraged to run using a bottle brush bristle and a shock grid at the end of the treadmill per the animal ethics protocol.

### 1.1.3 Micro-CT (µCT) imaging and analysis

Knee joints were fixed by immersion in 10% neutral buffered formalin at room temperature for 24 hours, prior to storage in 70% ethanol. On the day of scanning, all unstained samples were kept submerged in 70% ethanol and imaged with a 1172 SkyScan µCT at 4.83 µm resolution using a source voltage of 59LJkV, a source current of 167 µA, and an Al 0.5 mm energy filter. A rotation range of 180° with a rotation step of 0.5° and frame averaging of 2 was applied. Positioning of the samples during scanning were standardized, with the distal femur superior and the proximal tibia inferior in the scan view. Subsequent cartilage-staining with phosphotungstic acid (PTA) dissolved in 70% ethanol was then conducted on a subset of knee samples and imaged with the same parameters and protocol as stated above. Afterwards, all scans were reconstructed in NRecon software (v.1.7.0.4, Bruker microCT) using a smoothing width of 3, a beam-hardening correction of 100%, and a ring artifact correction of 20. Nomenclature used follows the recommendation of the American Society for Bone and Mineral Research. [17]

Pre-segmentation of the cartilage was conducted through Dragonfly (v.2022.1, Object Research Systems) and final segmentation was accomplished with the Biomedisa application (v.23.08.1, Australian National University CTLab) to measure cartilage thickness. Additionally, the subchondral bone was evaluated through Bone Analysis Wizard in Dragonfly to obtain average trabecular separation (Tb.Sp), average trabecular thickness (Tb.Th), trabecular number (Tb.N), and bone volume fraction (BV/TV). The optimal segmentation thresholds were automatically selected through the Otsu algorithm built into the Dragonfly software.

### 1.1.4 Histopathological assessment of the knee joint

At the experimental endpoint, a separate set of knee joints were fixed in 4% paraformaldehyde at room temperature for 24 hours on a shaker and then decalcified in 5% EDTA for 12 days at room temperature. Knees were processed and embedded in the sagittal plane in paraffin, and 6-μm– thick sections were cut from front to back. Sections were stained with toluidine blue. Three sections from the medial and lateral compartments were scored by 2 blinded reviewers using the Osteoarthritis Research Society International (OARSI) recommendations for histological assessments of OA in the mouse [18]. The average max score of each sample was then used for statistical analysis.

### 1.1.5 Synovial tissue pathological assessment

Considering the severity and bimodal distribution of the phenotype in these animals, we opted to take a more descriptive approach when describing the synovial changes that were occurring within these animals versus a semi-quantitative analysis. A pathologist with experience describing joint disease in animal models investigated the occurrence of specific pathological features observed in the synovium of the animals. The pathologist was blinded to all genotypes, sex, and activity. One hematoxylin and eosin (H&E)-stained section of the medial load-bearing region per animal was selected for analysis.

### 1.1.6 Histopathological assessment of the lumbar intervertebral discs

Lumbar spines were fixed for 24 hours with 4% paraformaldehyde, followed by 7 days of decalcification with Shandon’s TBD-2 (Thermo Fisher Scientific, Waltham, MA, USA) at room temperature. Tissues were embedded in paraffin and sectioned in the sagittal plane at a thickness of 5 μm. Mid-sagittal sections were deparaffinized and rehydrated as previously described [19] and stained using 0.1% Safranin-O/0.05% Fast Green. Sections were imaged on a Leica DM1000 microscope, with Leica Application Suite (Leica Microsystems: Wetzlar, DEU). To evaluate IVD degeneration, spine sections were scored by two observers blinded to age, exercise, sex, and genotype using a previously established histopathological scoring system for mouse IVDs [20]. Alterations to the scoring system were made, in which we excluded a score of 4 for the NP: “mineralized matrix in the NP”. To report on degeneration across the lumbar spine, scores for individual lumbar IVDs (L2-L6) were averaged, and the total score plotted for each individual mouse.

### 1.1.7 Enthesis and patellar tendon analysis

Mid-tendon sections were stained with toluidine blue and scored for distal quadriceps tendon enthesitis, distal patellar tendon enthesitis and patellar tendinopathy. Enthesitis was scored using the following parameters: 0 = normal; 1 = ectopic cartilage; 2 = ectopic cartilage/bone; 3 = ectopic cartilage/bone with a marrow cavity; 4 = ectopic bone with a marrow. Tendinopathy was scored using the following system: 0 = normal 1 = increased cellularity 2 = cell rounding/clustering 3 = chondrogenesis (i.e., proteoglycan-rich matrix, hypertrophy) 4 = ectopic cartilage/bone.

### 1.1.8 Statistics

The Department of Epidemiology and Biostatistics at Western University was consulted to determine the appropriate statistical analysis. Data are presented as stated in the respective figure. Prism 9 (GraphPad Software Inc.) Version 9.4.1 (458) was used to run all statistical tests including one-way analysis of variance (ANOVA) or two-way and three-way ANOVA for comparison. For OARSI scores, histopathological scores of the IVD, enthesitis and tendinopathy scores, males and females were analyzed separately within their respective genotypes and activity group. A Kruskal-Wallis test was used with an uncorrected Dunn’s test for multiple comparisons to determine statistical differences among the groups. Correlations were performed using Pearson’s r. All applicable data met assumptions for homoscedasticity or normality of residuals. Based on the recommendations of the editorial entitled: Moving to a World Beyond “p < 0.05” [21] the data is referred to in terms of weak (p < 0.05), moderate (p < 0.01) or strong (p < 0.001) statistical evidence. Please see Supplementary Tables 1-5.

## 1.2 Results

### 1.2.1 *Panx3* KO mice develop full-thickness cartilage erosion of the tibia and femur surface in aging

We first assessed whether *Panx3* KO mice develop histopathological OA compared to WT mice in aging, and whether forced treadmill running would influence these outcomes. Toluidine blue-stained, paraffin-embedded sections from the knee joints of 18-month-old WT and *Panx3* KO mice under SED or FEX conditions were analyzed (Figure 1A). We immediately noticed obvious osteoarthritic characteristics in the *Panx3* KO samples, including erosion of the cartilage and boney deposits in the enthesis (Figure 1A; Supplementary Figure 3,4). Consistent with our previous findings, closer histological assessment of the male mice observed full-thickness cartilage lesions in half of the *Panx3* KO mice in the medial tibial and femoral surfaces but not in any of the WT mice [13.15 mean rank difference, p = 0.0273; 10.33, p = 0.2661] (Figure 1B, C, D). In the lateral compartment, there was no statistical evidence for differences between SED WT and SED *Panx3* KO mice [4.011, p > 0.9999; 3.743, p > 0.9999] (Figure 1B, E, F). Regarding the effect of forced treadmill running, there was no statistical evidence that the addition of forced treadmill running influenced the cartilage structure in both knee compartments of WT mice for the tibia [8.200, p = 0.4613; 7.311, p = 0.6929] and the femur [0.5333, p > 0.9999; 0.7556, p > 0.9999]. In the lateral compartment, there was no statistical evidence that FEX *Panx3* KO mice had worse OARSI scores compared to SED WT mice, as 3 mice had full-thickness lesions of the tibia [11.80, p = 0.0979] (Figure 1E); and there was no statistical evidence for differences in femoral OARSI scores among the groups [5.419, p > 0.9999] (Figure 1F). Taken together, this suggests that some male *Panx3* KO mice develop histological features of severe OA only in the medial compartment of the tibia (approximately 50% phenotypic penetrance).

**Figure 1.**
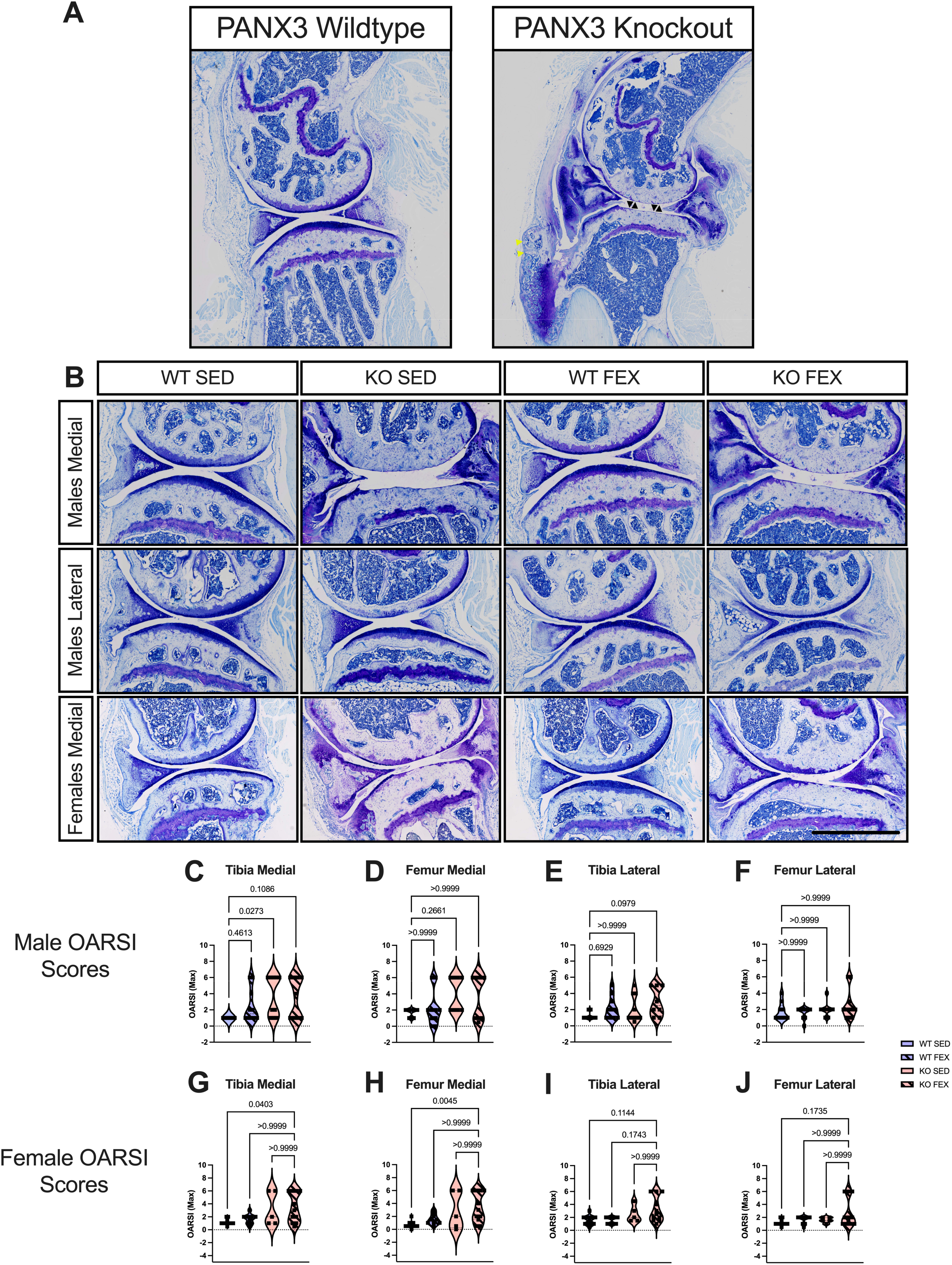
Male and female PANX3 KO mice develop full-thickness cartilage erosion in the tibia and femur in aging. Representative whole joint sagittal images stained with toluidine blue were taken at 10x magnification (scale bar = 1000 μm) at the medial and lateral compartments (A-B). (A) Full-view comparison of a PANX3 wildtype (WT) versus knockout (KO) joint. (B) Representative images of cartilage from male and female PANX3 WT and KO mice under sedentary (SED) and forced treadmill running (FEX) conditions, as indicated. (C-J) Violin plots delineating the distribution/grouping of OARSI max scores for male medial tibia (C) and femur (D), male lateral tibia (E) and femur (F), female medial tibia (G) and femur (H), and female lateral tibia (I) and femur (J). WT SED (N = 10 males, N = 12 females), WT FEX (N = 10 males, N = 9 females), KO SED (N = 10 males, N = 5 females), KO FEX (N = 9 males, N = 11 females). For statistical comparisons among the groups, a Kruskal-Wallis test followed by a Dunn’s multiple comparisons test was performed. Black arrows point to full thickness cartilage loss, yellow arrows point to boney deposits in the enthesis.

Next, we performed the same histological analysis of female knees from aged WT and *Panx3* KO mice under SED and FEX conditions. Note, female mice had not been included in our earlier aging study [13]. Like in males, several female *Panx3* KO mice developed full-thickness cartilage erosion in the medial compartment (6 mice in total), while no WT mice developed such erosion [11.79, p = 0.0403; 14.95, p = 0.0045] (Figure 1B, G, H). Forced treadmill running seemed to have little to no effect on cartilage structure in the medial compartment [1.700, p > 0.9999 ; 5.255, p > 0.9999] (Figure 1B, G, H). In the lateral compartment, while some SED *Panx*3 KO mice showed signs of cartilage erosion, full thickness erosion was only observed in some *Panx3* KO mice that were forced to run (N = 3), but there was no statistical inference to suggest differences between these groups (Figure 1I, J). This data suggests, that like male *Panx3* KO mice, female mice exhibit a bimodal distribution, with a subset of mice developing full-thickness cartilage erosion, and the addition of forced treadmill running may influence development of cartilage loss of the tibia in the lateral compartment with approximately 50% phenotypic penetrance.

Given we saw full-thickness lesions of the tibia, we conducted further μCT analysis on PTA-stained cartilage samples with the contralateral knee of the WT and *Panx3* KO. In support of our histopathological observations, all *Panx3* KO mice demonstrated moderate statistical evidence for reduced tibial cartilage thickness in comparison to WT mice [F (1, 22) = 8.83, p = 0.0070], regardless of sex and activity (Figure 2A, B). This data suggests that both *Panx3* KO male and female mice experiences cartilage erosion throughout the whole cartilage of the tibia even without forced exercise as no statistical effect was observed between activity groups. In comparison to our previous work [38], this may suggest the development of cartilage loss of the tibia is due to a combination of both natural aging and genotype.

**Figure 2.**
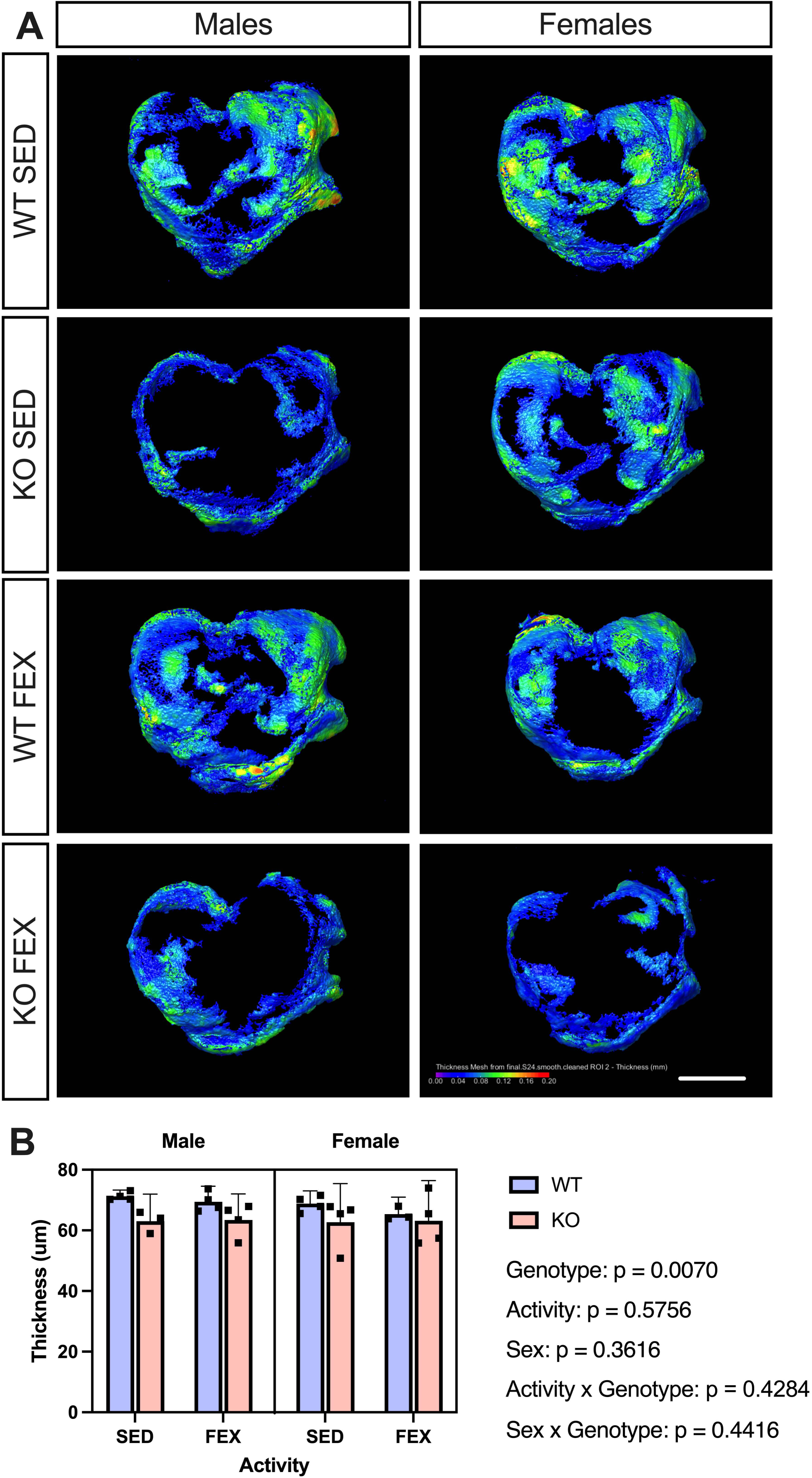
PANX3 KO mice have reduced cartilage thickness when examined through micro-CT. Representative micro-CT images of knee cartilage samples stained with phosphotungstic acid from male (left panel) and female (right panel) PANX3 WT and KO mice subjected to sedentary (SED) or forced treadmill running (FEX) conditions, as indicated (A). Images shown from the superior view (scale bar = 1 mm). (B) Corresponding micro-CT cartilage thickness measurements of treatment groups WT SED (N = 4 males, N = 4 females), WT FEX (N = 4 males, N = 3 females), KO SED (N = 3 males, N = 4 females), KO FEX (N = 4 males, N = 4 females). For statistical comparisons among the groups, a three-way ANOVA with Sex x Genotype x Activity as factors followed by Tukey’s multiple comparisons test was performed. All data are shown as means ± CI.

### 1.2.2 *Panx3* KO mice exhibit reduced trabecular thickness and secondary ossification center volume in the proximal tibia

Building on the established findings regarding the pivotal role that PANX3 plays in the regulation of osteoblast differentiation, mechanically induced bone modelling, and bone formation [3, 23, 24], our prior observations in 24-30-week-old mice revealed reductions in the bone volume fraction of the proximal tibia [38]. Considering these insights, and the significant articular cartilage degeneration we are witnessing in the current study, we next wanted to identify any pathological changes in the secondary ossification center of the proximal tibia through μCT imaging (Figure 3A). *Panx3* KO mice were observed to have weak statistical evidence of an increase in average trabecular separation [F (1, 71) = 5.53, p = 0.0215], which is representative of cavities in the bone marrow (Figure 3A, B), and a complementary decrease in average trabecular thickness [F (1, 71) = 4.11, p = 0.0462] (Figure 3A, C). Additionally, there was weak statistical evidence for a two-way interaction of activity x genotype [F (1, 71) = 5.87, p = 0.0180], where the presence of forced treadmill running seemed to reduce the genotypic-increase in average trabecular separation of the male *Panx3* KO mice and completely reverse its affects in the female mice (Figure 3B). However, there was no evidence of any genotypic or activity changes to trabecular number [F (1, 71) = 2.56, p = 0.1141] (Figure 3D). Despite this, further analysis of the tibial secondary ossification center revealed strong evidence for a genotypic effect on bone volume fraction [F (1, 71) = 14.1, p = 0.0003], with *Panx3* KO mice having reduced bone volume compared to WT mice (Figure 3E). There was also weak statistical evidence for an interaction between activity and genotype [F (1, 71) = 4.58, p = 0.0358], and moderate statistical evidence for a sex x genotype interaction [F (1, 71) = 10.6, p = 0.0017]. More specifically, a sedentary lifestyle seems to exacerbate the bone volume reduction observed in *Panx3* KO mice, whereas this bone volume decrease also appears to be absent in females compared to males (Figure 3E). Altogether, this data suggests that a *Panx3* deletion leads to a reduction in trabecular bone within the secondary ossification center of the proximal tibia and the presence of forced exercise may mitigate the extent of this alteration.

**Figure 3.**
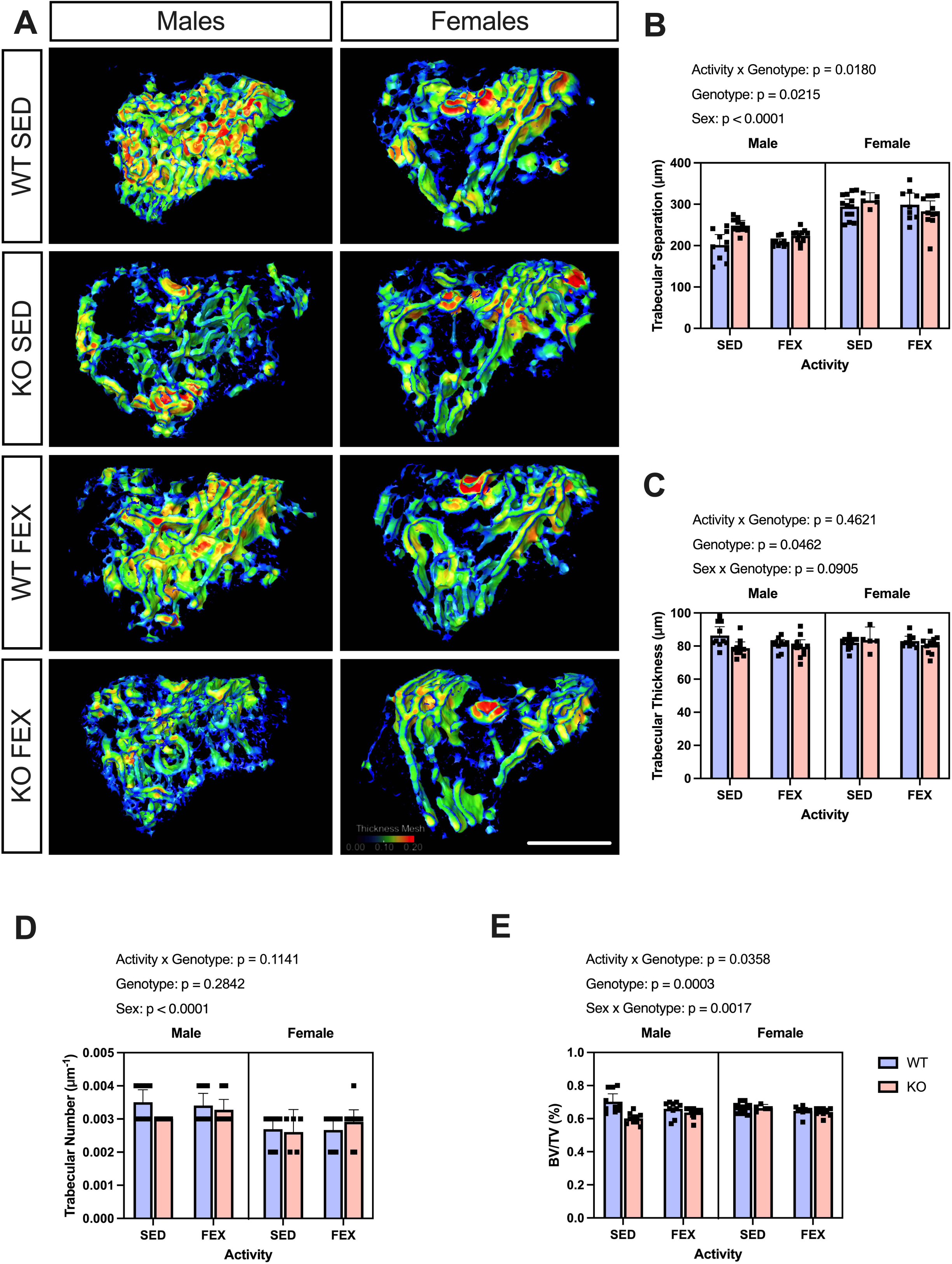
A PANX3 deficiency alters subchondral bone characteristics in aged mice. Representative micro-CT 3D renderings of male (left panel) or female (right panel) mouse knee subchondral bone samples from indicated genotype and treatment group (A). Scale bar = 1 mm. (B-E) Bar graphs displaying the average trabecular separation (B), average trabecular bone thickness (C), average trabecular number (D), and subchondral bone volume fraction (E). WT SED (N = 10 males, N = 13 females), WT FEX (N = 10 males, N = 9 females), KO SED (N = 10 males, N = 5 females), KO FEX (N = 11 males, N = 11 females). For statistical comparisons among the groups, a three-way ANOVA with Sex x Genotype x Activity as factors followed by Tukey’s multiple comparisons test was performed. All data are shown as means ± CI.

### 1.2.3 *Panx3* KO mice have comparable body weights to WT mice under sedentary and forced treadmill running conditions within their respective sex

Having previously published large reductions in body weights of *Panx3* KO mice compared to WT mice in adulthood [22], we next wanted to investigate whether the observed changes in articular cartilage and trabecular bone were attributable to alterations in mechanical loading that stemmed from differences in body weight between the genotypes. Hence, we also investigated the effect of deleting *Panx3* on body weight in aged animals (Supplementary Figure 1) and found that those genotypic differences were diminished with age. There was no statistical evidence for differences in body weight between genotypes in both male [F (1, 36) = 0.2273, p = 0.6364] (Supplementary Figure 1A) and female [F (1, 32) = 3.207, p = 0.0828] (Supplementary Figure 1B) mice. While there was no statistical evidence for body weight differences between activity groups in males [F (1, 36) = 0.09, p = 0.7574], there was weak statistical evidence for female mice to have lower body weights when forced to run on a treadmill for 6 weeks compared to SED female mice [F (1, 32) = 5.37, p = 0.0270]. This data suggests aged WT and *Panx3* KO mice have similar body weights under SED and FEX conditions, thereby suggesting body weight differences are unlikely to be a contributing factor for the observed genotypic disparities in the joint tissue.

### 1.2.4 *Panx3* KO mice develop mild to severe synovitis under both sedentary and forced treadmill running conditions

Considering the bimodal distribution seen in our KO mice, and upon observation of the severe synovial changes in the mice with full-thickness erosions, we chose to take a descriptive approach for the synovial analysis. Synovial tissue was assessed by a blinded pathologist with extensive experience describing animal model synovial tissue. H&E-stained sections in the medial load-bearing zone were assessed. Three distinct joint morphological phenotypes were observed across the groups: 1) no lesions (Figure 4A); 2) mild acute synovitis (Figure 4B); and 3) severe diffuse synovial fibrosis with ectopic ossification (Figure 4C). All samples with mild to severe synovitis were from *Panx3* KO mice, except for one WT sample. Synovium of *Panx3* KO mice with intact cartilage consisted of acute lymphocytic synovitis, where the synovium was expanded by lymphocytes and few macrophages (Figure 4B mild synovitis). Full-thickness cartilage erosion in the *Panx3* KO mice coincided with severe diffuse synovial fibrosis, ulceration, and ectopic ossification (Figure 4C severe synovitis). These mice had locally extensive to complete effacement of the synovium by collagen, fibrocartilage and in some cases, bone interrupted by areas of acellular basophilic material. These findings suggest that aged *Panx3* KO mice develop mild-to-severe synovitis of the knee joint, which coincides with the severity of cartilage erosion.

**Figure 4:**
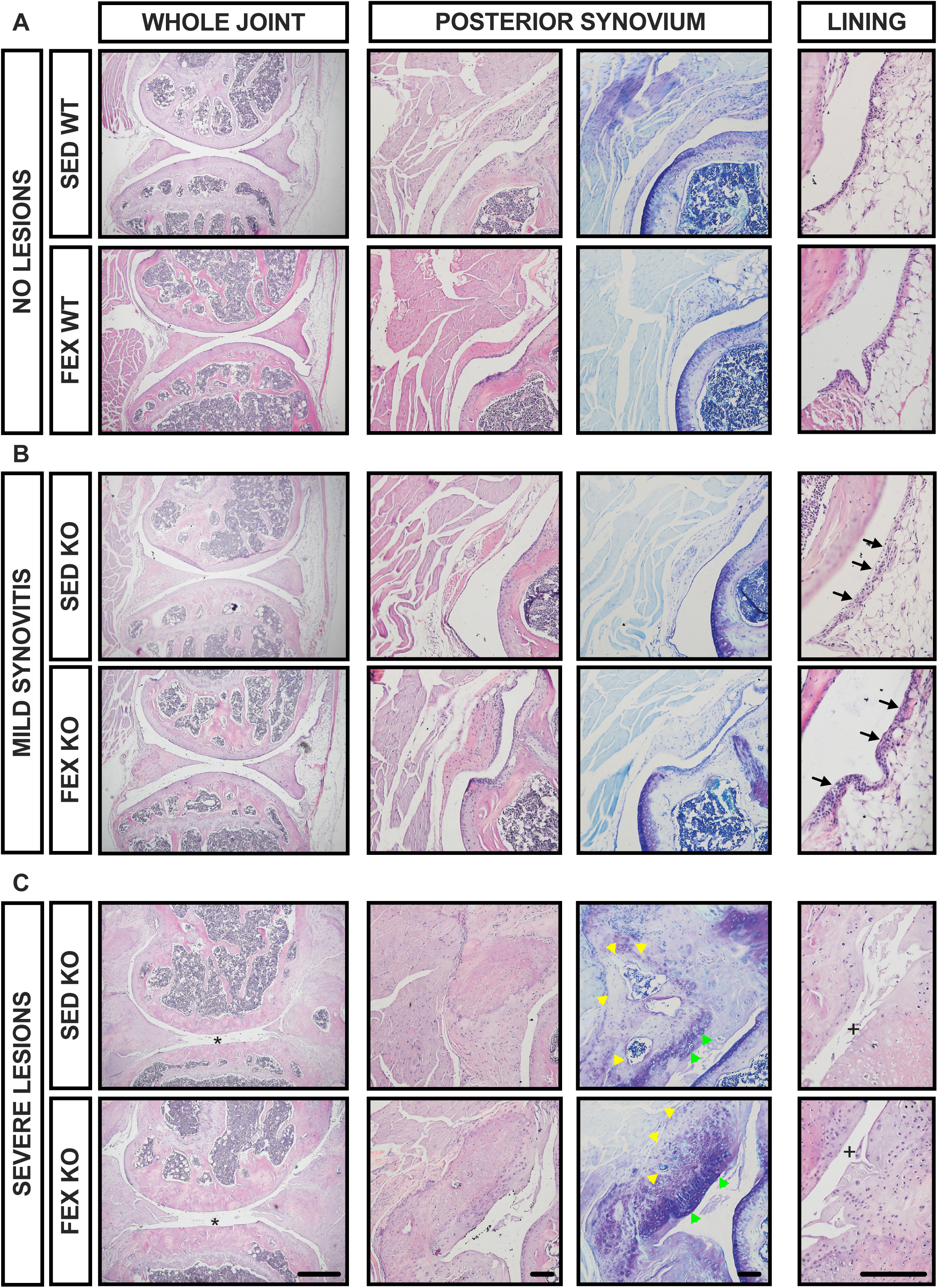
*Panx3* KO mice develop mild to severe synovitis of the knee in aging. Sagittal sections of the medial compartment were stained with H&E and assessed by a blinded pathologist. Slides were assessed for signs of immune cell infiltration of the synovial lining (black arrows), indicating mild synovitis, which was a phenotype of *Panx3* KO mice (one WT mouse was characterized to have mild synovitis). Ectopic ossification (yellow arrows) and fibrocartilage (green arrows) were observed exclusively in *Panx3* KO mice and was characterized as severe lesions. * Denotes cartilage erosion. + denoting loss of synovial lining. Whole joint images (Left) are 4x magnification. Scale bar = 500μm. Posterior synovium (middle) are 10x magnification. Scale bar = 100μm. Lining (right) images are 20x magnification. Scale bar = 100μm.

### 1.2.5 *Panx3* KO mouse IVDs age normally in males and females even when forced to treadmill run

To investigate the role of PANX3 in age-associated IVD degeneration, and whether forced treadmill running in aging influences disc health, we next analyzed the IVDs for histological changes. WT and *Panx3* KO mice were aged to 18 months and lumbar spines were analyzed histologically as described in the Methods section. Under both SED and FEX conditions, male (Supplementary Figure 2A, B) and female (Supplementary Figure 2C, D) *Panx3* KO IVDs appeared normal relative to WT controls with no differences in histopathological features of degeneration in the nucleus pulposus (NP) and the AF at any disc height (Supplementary Figure 2A, C) or when averaged across the lumbar spine (Supplementary Figure 2B, D). This data suggests that the IVDs of both male and female *Panx3* KO mice, age and respond to forced treadmill running similarly to their WT counterparts as no effect was observed under the applied conditions.

### 1.2.6 Male *Panx3* KO mice display ectopic cartilage and bone deposits containing marrow in the enthesis of the quadriceps and patellar tendons

Considering the enthesis originates from fibrocartilage cells that are highly responsive to mechanical loading [23], we next analyzed the patellar tendon for signs of tendinopathy and enthesitis at the distal patella and quadriceps tendons (Figures 5&6A). In male mice, there was weak statistical evidence [12.26, p = 0.0450] suggesting SED *Panx3* KO mice develop enthesitis of the quadriceps tendon enthesis, consisting of cartilage and bone deposits, often with a marrow cavity (Figure 5B). Throughout the patellar tendon, there was no statistical evidence for histopathological cellular changes in any of the groups [p = 0.1473] (Figure 5C). Like the quadriceps enthesis, the distal patellar tendon enthesis of SED *Panx3* KO mice showed signs of enthesitis, including deposits of cartilage and bone, often with a marrow cavity, compared to SED WT mice [11.31, p = 0.0205] (Figure 5D). In female mice, we performed the same semi-quantitative analysis of the enthesis and patellar tendon (Figure 6A). At the quadriceps enthesis, there was weak statistical evidence [14.50, p = 0.0246] that *Panx3* KO mice develop enthesitis with forced treadmill running (Figure 6B), with all the mice showing ectopic cartilage and bone, often with marrow formation, within the quadriceps tendon enthesis (Figure 6E). Throughout the patellar tendon, there was no statistical evidence [p = 0.4007] for histopathological changes to cellular shape or distribution between any of the groups (Figure 6C). At the patellar tendon enthesis, there was weak statistical evidence [8.063, p = 0.0201] that WT mice develop enthesitis with forced treadmill running, with no statistical evidence of histological differences in *Panx3* KO mice (Figure 6D). This data suggests that *Panx3* KO mice develop enthesitis of the quadriceps and patellar tendon entheses during aging, while forced treadmill running in female WT mice produced enthesitis at the distal patellar tendon enthesis. We next ran a Pearson’s correlation between medial OA scores and the enthesitis and tendinopathy scores to determine if there was a relationship between which mice develop OA and those that develop enthesis and tendon pathology. There was strong statistical evidence suggesting that higher enthesitis and tendinopathy scores coincided with higher OA scores (Table 1).

**Figure 5:**
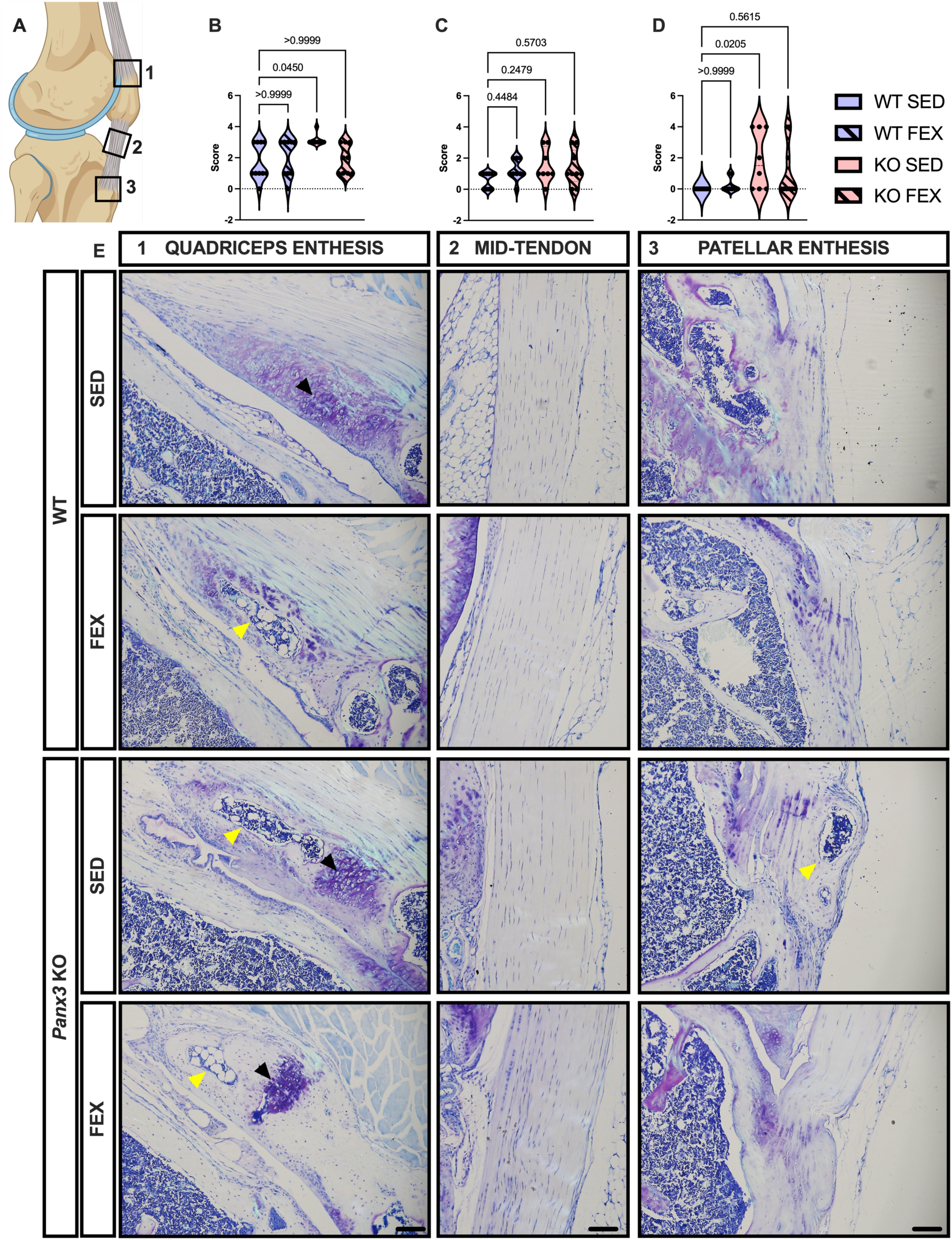
Sedentary male *Panx3* KO mice show signs of patellar and quadriceps enthesitis. Male WT and *Panx3* KO knee sections were stained with toluidine blue, sectioned in the sagittal plane and scored for quadriceps enthesitis (A1), patellar tendinopathy (A2), and patellar enthesitis (A3). Sedentary (SED), and forced treadmill running (FEX). Violin plots showing distribution/grouping of histological scores for quadriceps enthesitis (B), tendinopathy (C), and patellar enthesitis (D). Representative toluidine blue sagittal sections (E). 10x magnification. Scale bar = 100 μm. Black arrows point to cartilage, yellow arrows point to bone and marrow. WT SED (N = 8), WT FEX (N = 10), KO SED (N = 12), and KO FEX (N = 11). For statistical comparisons among the groups, a Kruskal-Wallis test was performed.

**Figure 6:**
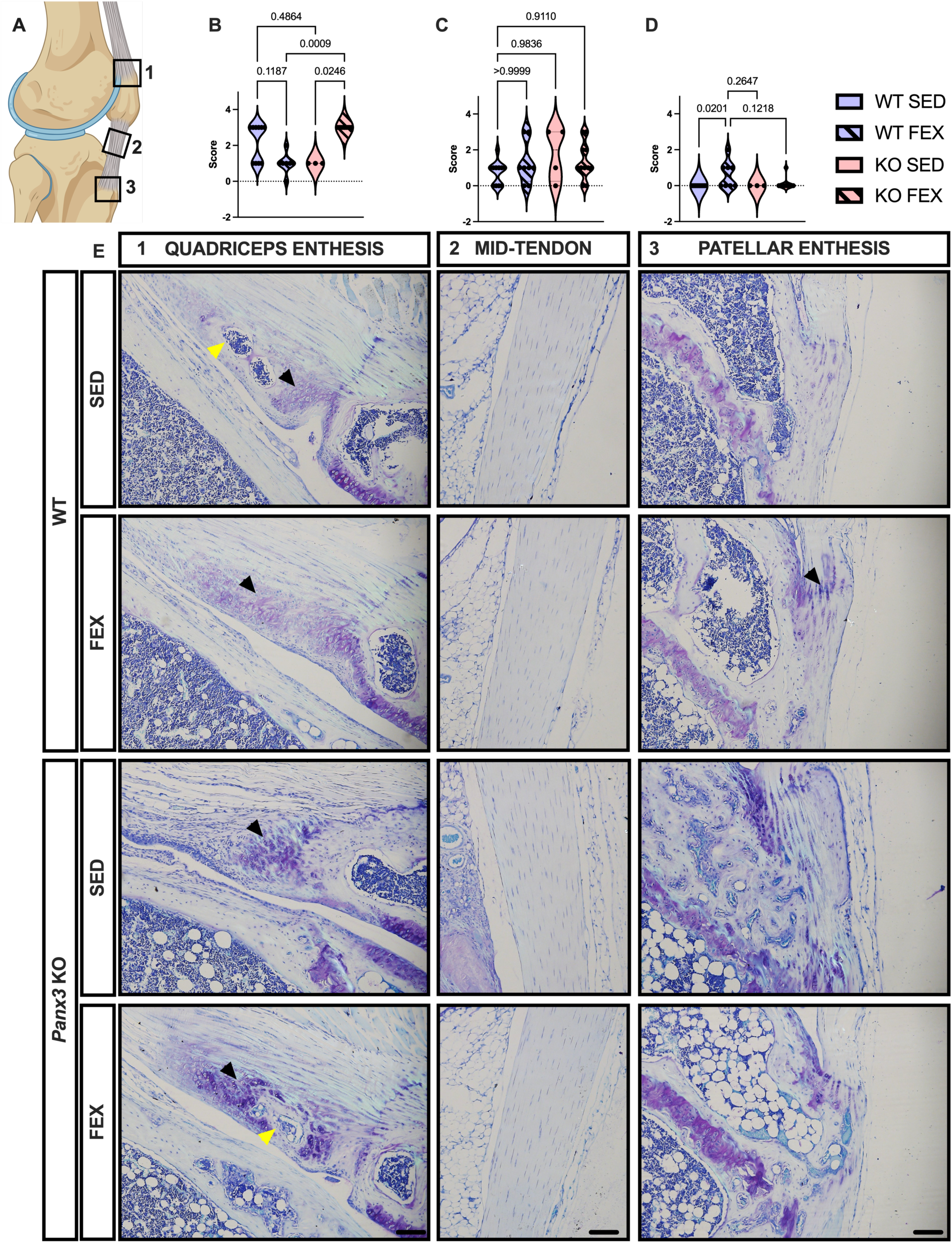
Female *Panx3* KO mice develop quadriceps enthesitis with forced treadmill running. Female WT and *Panx3* KO knee sections were stained with toluidine blue, sectioned in the sagittal plane and scored for quadriceps enthesitis (A1), patellar tendinopathy (A2), and patellar enthesitis (A3). Sedentary (SED), and forced treadmill running (FEX). Violin plots showing distribution/grouping of histological scores for quadriceps enthesitis (B), tendinopathy (C), and patellar enthesitis (D). Representative toluidine blue sagittal sections (E). 10x magnification. Scale bar = 100 μm. Black arrows point to cartilage, yellow arrows point to bone and marrow. WT SED (N = 14), WT FEX (N = 8), KO SED (N = 4), and KO FEX (N = 11). For statistical comparisons among the groups, a Kruskal-Wallis test was performed.

**Table 1:** Medial tibial OARSI scores correlate with tendinopathy and enthesitis scores.

|  | OARSI Score | Tendinopathy | Quadriceps Enthesitis | Patellar Enthesitis |
| --- | --- | --- | --- | --- |
| OARSI Score | - |  |  |  |
| Tendinopathy | 0.38 $p = 0.001$ | - | | |
| Quadriceps Enthesitis | 0.38 $p = 0.002$ | 0.27 $p = 0.029$ | - | |
| Patellar Enthesitis | 0.58 $p < 0.001$ | 0.32 $p = 0.008$ | 0.19 $p = 0.127$ | - |

## 1.3 Discussion

In this study, we found that 18-month-old male and female *Panx3* KO mice demonstrate severe OA of the knee and WT control mice demonstrated milder OA, while IVDs seem to be histologically comparable between genotypic groups. The addition of forced mechanical loading through treadmill running does not exacerbate this phenotype in the medial compartment, while a subset of FEX *Panx3* KO mice had full-thickness lesions in the lateral compartment, suggesting a compartment-specific effect of forced treadmill running in *Panx3* KO mice. The presence of forced exercise also attenuated the increase in average trabecular separation and decrease in bone volume fraction of the proximal tibial secondary ossification center for *Panx3* KO mice in comparison to WT. This observation may result from increased bone remodelling triggered by mechanical stress, where the bone adapts to increased loading by minimizing the space between the trabeculae to strengthen its framework. However, due to the male *Panx3* KO mice already having a more osteopenic phenotype, and hence a greater trabecular separation and lower bone volume fraction, this interaction manifests as less pronounced changes in the separation or bone volume between the genotypes. It is notable that in the histology we witnessed the presence of blood entering the joint cavity, which may be an indication that minor trauma or stresses on the bone can result in a fracture that causes bleeding into the joint space. Additionally, a fracture may also contribute to the development of osteoarthritis overtime by altering the biomechanics and stability of the joint. Additionally, *Panx3* KO mice develop mild to severe synovitis consisting of lymphocyte infiltration, and ectopic fibrocartilage and calcification of the knee joint. Male *Panx3* KO mice also appeared to have histological features of quadriceps and patellar tendon enthesitis under SED conditions, whereas female *Panx3* KO mice developed quadriceps tendon enthesitis with forced treadmill running. Within lumbar spine IVDs, both male and female *Panx3* KO mice had similar histopathological features compared to the WT controls. Even with the stress of forced treadmill running, there was no statistical evidence for histopathological differences among the groups, suggesting running later in life is not detrimental to disc structure in either genotype.

Full-thickness cartilage erosion is not a normal histological feature of knee joints in aged mice [24]. In this study, the full-thickness cartilage loss observed in our animals was accompanied by erosion and fibrillation of adjacent cartilage surfaces. Additionally, using μ-CT contrast imaging of cartilage in the contralateral knee, we found that these mice had reduced thickness in the *Panx3* KO mice. However, it is notable that the representative images show full cartilage loss in both the WT and KO mice, albeit more in mice with the latter genotype. But given that none of the WT mice had full-thickness cartilage loss in the histological analysis, this supports that the ulcerations we see may be artifacts from processing (i.e., ineffective PTA-staining of degenerative areas) and thereby making it hard to determine if there are the presence of pre-mortem full-cartilage lesions. All this is not to suggest that the cartilage of these *Panx3* KO mice is not degenerating, but rather histological and μ-CT representation may not accurately depict the *in vivo* state of the cartilage tissue pre-mortem.

In addition to cartilage erosion, we saw mild to severe synovitis in the *Panx3* KO animal knees, which included extensive fibrocartilage and calcification deposition within the synovium. Given this phenotype was not detected in WT mice, we suspect that it may be *Panx3* KO specific. Whether PANX3 is expressed in synovial tissue has not been determined; however, *in silico* data suggests *PANX3* should be expressed in human synovial fibroblasts [25]. Additionally, considering the severe inflammatory phenotype, we are unaware of any indication that macrophages or lymphocytes express *Panx3.* Future studies should investigate the periarticular expression and function of *Panx3* in joint tissues.

Considering the evidence showing PANX3’s role in cartilage, it is possible that the synovial phenotype is initiated by cartilage degradation and subsequent synovitis. Chronic release of damage-associated molecular patterns or other catabolic signals (e.g. cytokines) from degrading cartilage into the synovial fluid space may activate synovial lining macrophages [26]. In our previous report challenging 30-week-old *Panx3* KO mice, we showed superficial cartilage erosion which was exacerbated by forced treadmill running and resulted in moderate evidence of synovial lining thickening, suggesting early OA development [38] A lifetime of cartilage erosion may be chronically stimulating synovial macrophages leading to these pathological changes.

Previously, we found that aged *Panx3* KO mice had low lubricin expression in the superficial zone of the articular cartilage [13]. Lubricin is an essential lubricating protein for the joint surface [27], and *in vitro* models have shown that lubricin has anti-inflammatory effects on synovial lining fibroblasts by binding to toll-like receptors 2 and 4 [28]. Taken together, the superficial erosion in adulthood of *Panx3* KO mice may lead to reduced lubricin levels in aging, which could be chronically activating synovial lining cells, and thus producing the severity of synovitis we observed in the present study.

Inflammation is associated with age-related pathologies [29] including primary OA, and nuclear Factor Kappa β (NF-κβ) is a proposed central pathway of inflammation in OA [24]. Interestingly, through mechanotransduction pathways, chondrocytes can release ATP, and this extracellular ATP has been shown to activate NF-κβ signalling and contribute to OA [30, 31]. While some ATP release is required to maintain normal cartilage homeostasis [32], abnormal mechanical loading of cartilage increases chondrocyte ATP release [33, 34]. This suggests that there are physiologically healthy levels of ATP release required for cartilage maintenance, but dysregulation of this mechanism could contribute to inflammation and OA. Our previous reports showed that aging WT mouse cartilage maintains similar PANX3 protein expression at 6, 18, and 24 months of age [13], suggesting PANX3 is required to maintain cartilage health well into aging. Deletion of *Panx3* may dysregulate this ATP signalling given its canonical function as a mechanosensitive, plasma membrane ATP release channel in cells such as chondrocytes [4, 35].

In our previous report, *Panx3* deletion did not significantly impact the progression of age-associated histopathological IVD degeneration in male mice at 18 and 24 months of age compared to WT mice [6]. The present study also determined that the aged female *Panx3* KO mice IVD histopathological analysis matched that of males. The contrasting difference between the knee joint and IVDs of *Panx3* KO mice is interesting considering the similar mechanism of disease progression between OA and IVD diseases [36]. It appears that PANX3 is not essential to IVD health during normal aging, as our previous report showed low transcript expression of *Panx3* in IVDs from 6 to 24 months of age relative to levels at 2 months of age [6]. It is possible that PANX3 is utilized in early life and development of the IVD, while dispensable in aging. Interestingly, forced treadmill running seemed to have no effect on histological features of the IVDs regardless of genotype or sex. This was surprising, considering *Panx3* KO mice developed histopathological features in the AF of IDD with forced treadmill running in younger adult mice [38]. It was reasonable to hypothesize that aging would have rendered these mice more susceptible to forced treadmill running-induced changes to the IVD. However, considering aging alone results in relatively severe spontaneous IDD in mice [37], any potential impact of forced treadmill running, positive or negative, may have been undetectable with histopathological scoring.

With all this in mind, we propose employing the aged *Panx3* KO mouse model utilized in this study as a novel model of primary knee OA. In contrast to many existing mouse models of spontaneous age-induced OA, which often exhibit high variability in disease incidence [39], this mouse model demonstrated a notable incidence rate of over 40% for severe OA. Moreover, the bimodal distribution of OARSI scores indicates that our *Panx3* KO model also displays low variation in disease severity. Additionally, this model only requires a single gene deletion and does not require the intervention of exercise protocols such as forced treadmill running, thereby reducing the technical demands associated with both its generation and maintenance. This minimization of demands also facilitates its integration with other models and interventions.

Future studies should involve a time course of *Panx3* KO mouse OA development. Specifically, analyzing earlier time points (for example. 6, 9, and 12 months) to determine when distinct OA phenotypes arise, and to characterize its progression. This will allow for the analysis of the early cellular changes that may be driving this severe OA. Additionally, considering that obesity is a strong risk factor for OA, and coincides with aging [2], high-fat diet studies in *Panx3* KO mice are also warranted. Lastly, tissue-specific KO models using various Cre driver lines are also warranted to determine the cell type responsible for this severe OA development.

## 1.4 Conclusion

Approximately half of aged male and female *Panx3* KO mice develop severe OA under both SED and FEX conditions, which occurs spontaneously as in human primary OA. Hence, we suggest the potential of the *Panx3* KO mice as a novel primary osteoarthritis model. Additionally, *Panx3* KO mice have enthesitis of the quadriceps and distal patella more often than WT mice, but at the lumbar IVDs, *Panx3* KO mice have similar histological features as WT mice in aging. Collectively, our data suggests chronic suppression of PANX3 throughout life may be contraindicated and cause severe OA. Potential therapeutic interventions could include agonists of PANX3 to overcome the lower expression or reduced functionality of PANX3 channels in aged joint tissues.

## CONFLICT OF INTEREST

The authors declare no competing or conflicting interests of any kind.

**Supplementary Figure 1: Aged *Panx3* KO mice have similar body weights to WT mice under sedentary and forced treadmill running conditions within each sex.**

WT and *Panx3* KO mice were aged to 18 months of age and randomized to either sedentary (SED) or forced treadmill running (FEX) for 6 weeks. Body weights were taken at termination. WT SED (N = 9), WT FEX (N = 10), KO SED (N = 11), and KO FEX (N = 10), as indicated. Male body weights (A). Female body weights (B). WT SED (N = 12), WT FEX (N = 9), KO SED (N = 4), and KO FEX (N = 11), as indicated. For statistical comparisons among the groups, a two-way ANOVA with genotype x activity was used with a Tukey’s correction for multiple comparisons. All data are shown as means ± CI.

**Supplementary Figure 2: No overt differences in histopathological scores of the IVDs between WT and *Panx3* KO mice.**

Wildtype (WT) and *Panx3* KO (KO) mouse lumbar spines were stained with Safranin O Fast Green. Histopathological scores for males by disc height (A) and average disc score across the lumbar spine (B). WT SED (N = 8), WT FEX (N = 10), KO SED (N = 12), and KO FEX (N = 11). Female histopathological scores by disc height (C) and average disc score across the lumbar spine (D). WT SED (N = 14), WT FEX (N = 8), KO SED (N = 4), and KO FEX (N = 11). For statistical comparisons among the groups, a Kruskal-Wallis test was performed. All data are shown as means ± CI.

**Supplementary Figure 3. Full-view histological image of male mice knee joints.**

Representative histological images of whole knee joints from wildtype (WT) or PANX3 knockout (KO) male mice that underwent sedentary (SED) or forced exercise (FEX) treatment. Samples were sagittally sectioned and stained in toluidine blue. Images were captured at 10x objective (scale bar = 1000 μm) in the medial and lateral compartments.

**Supplementary Figure 4. Full-view histological image of female mice knee joints.**

Representative histological images of whole knee joints from wildtype (WT) or PANX3 knockout (KO) female mice that underwent sedentary (SED) or forced exercise (FEX) treatment. Samples were sagittally sectioned and stained in toluidine blue. Images were captured at 10x objective (scale bar = 1000 μm) in the medial and lateral compartments.

