## Supplementary figures and images for "Severe Osteoarthritis in Aged PANX3 Knockout Mice: Implications for a Novel Primary Osteoarthritis Model"

### Suppl. 1-4

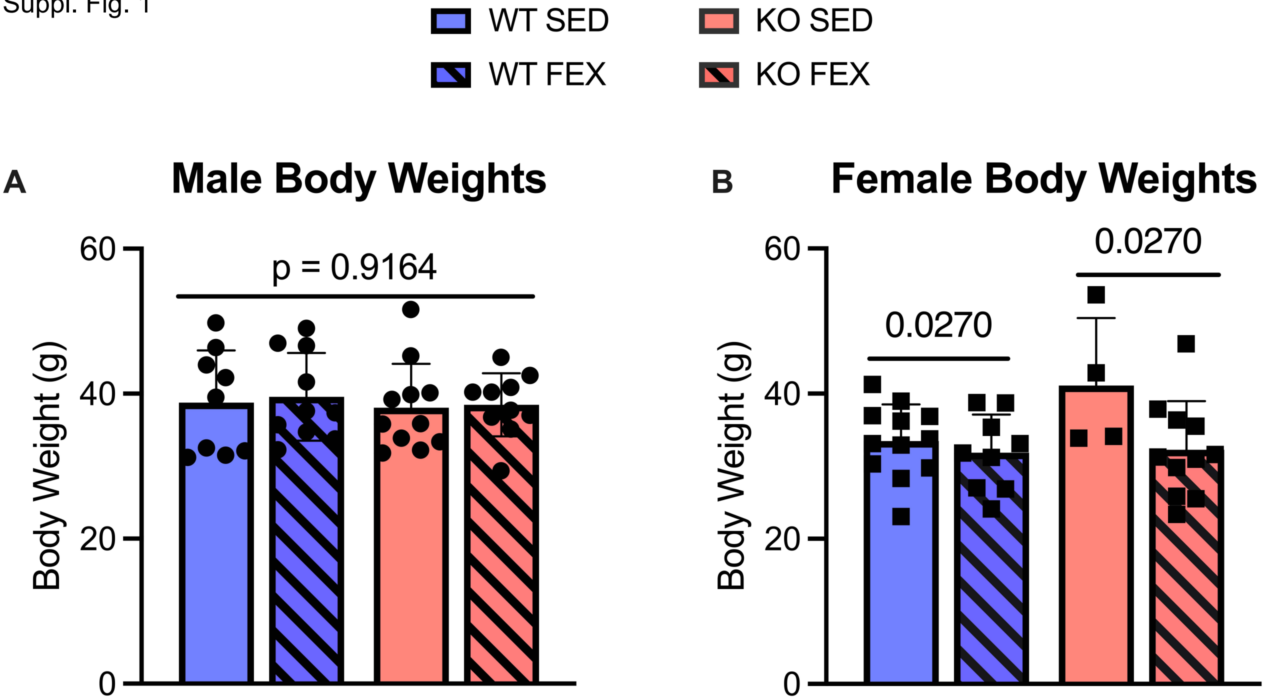

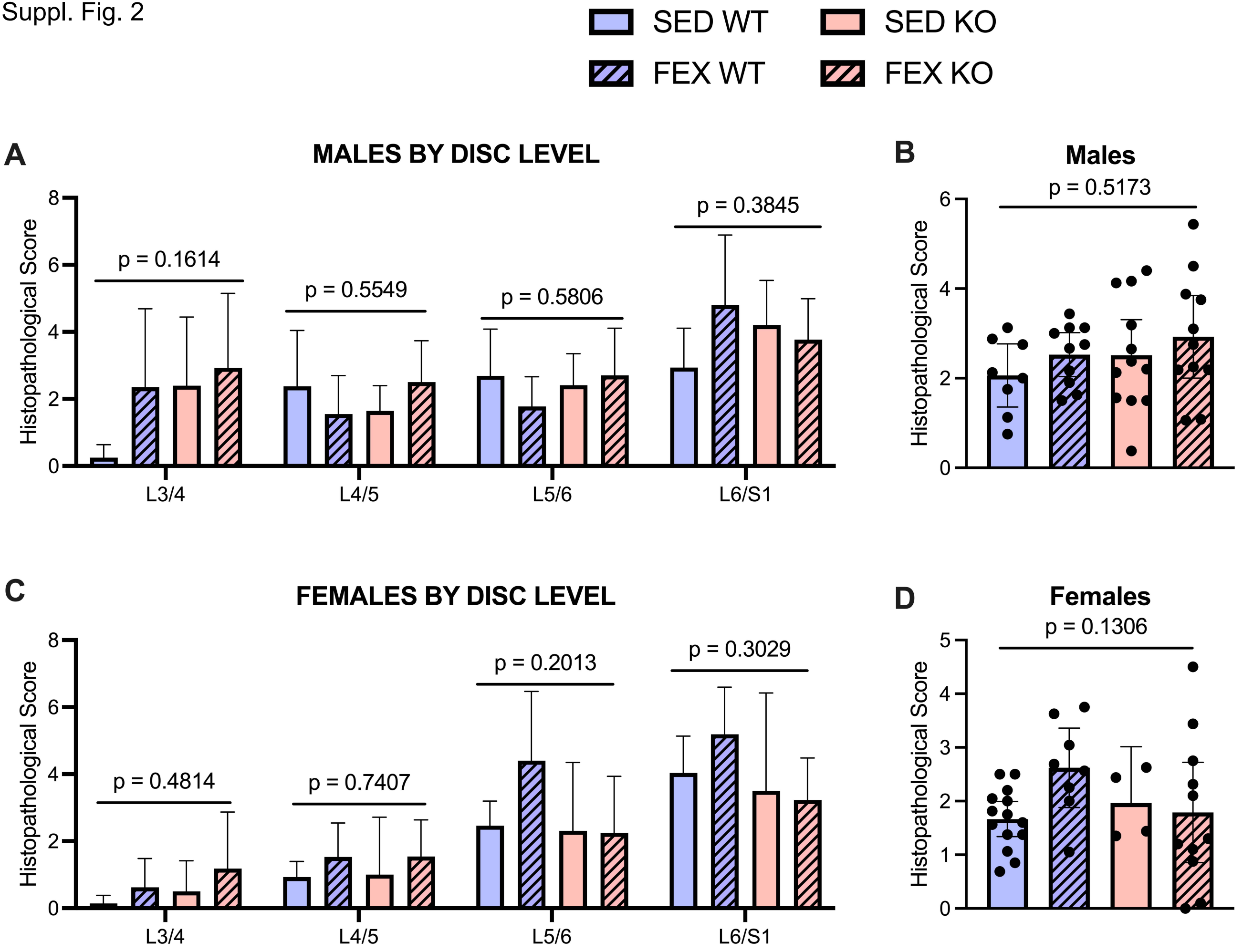

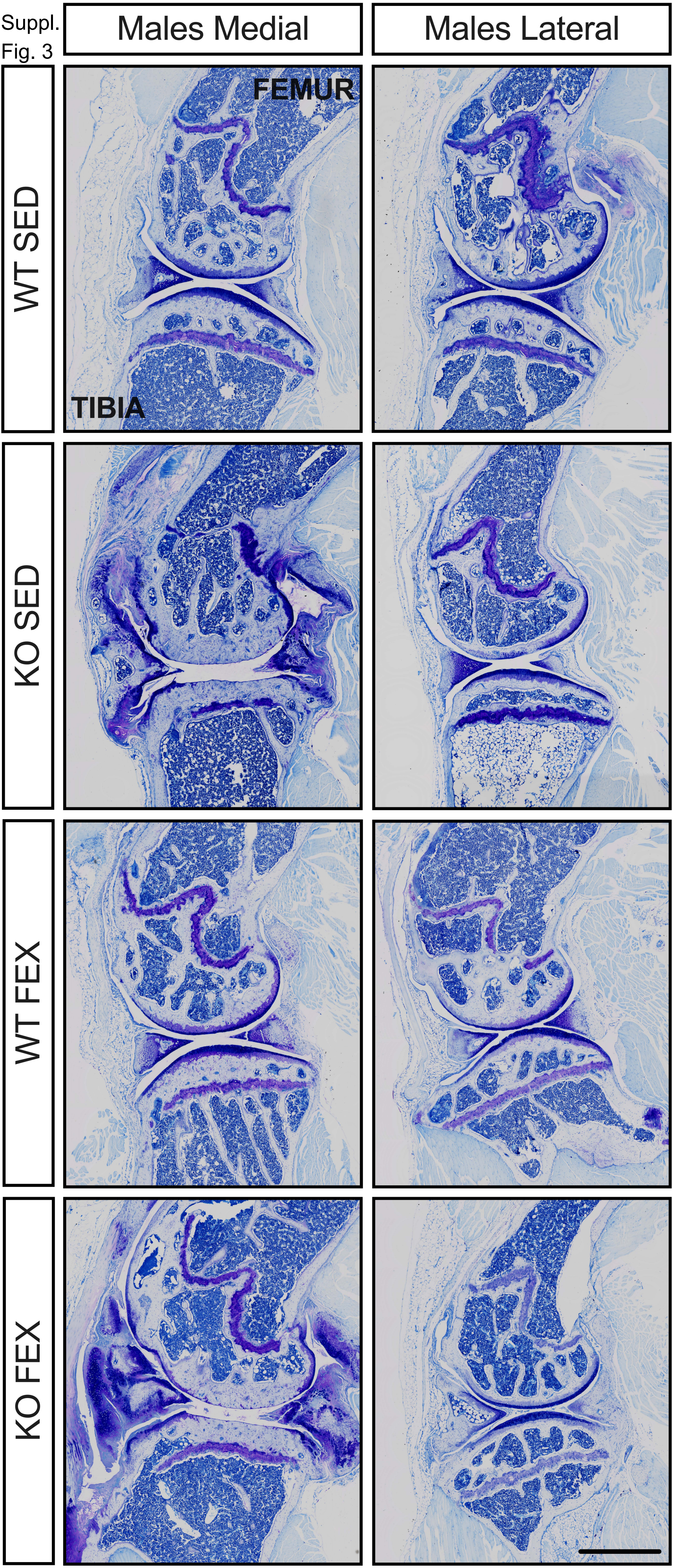

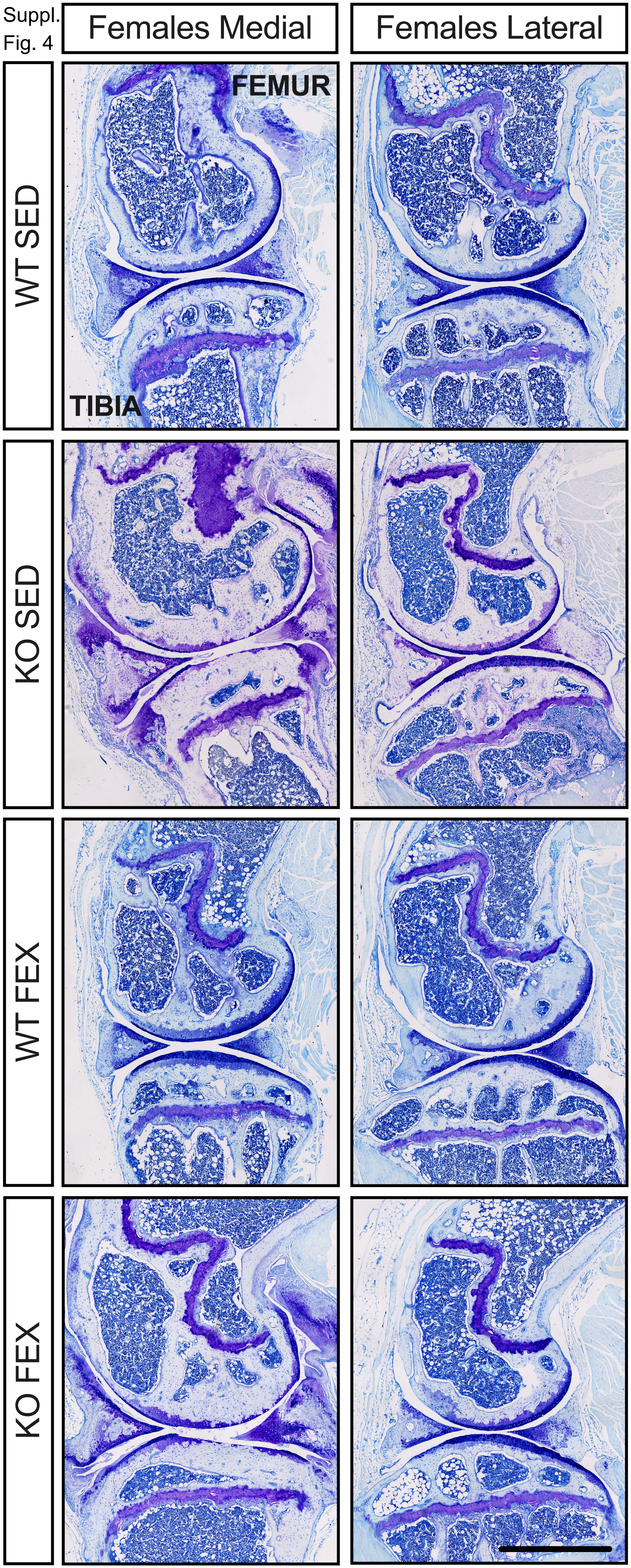
